# Aging concomitantly reduces skeletal muscle proteome plasticity and hypertrophic responses to resistance training

**DOI:** 10.64898/2026.04.27.721183

**Authors:** Dustyn T. Lewis, J. Max Michel, Mason C. McIntosh, Dakota R. Tiede, Daniel L. Plotkin, Madison L. Mattingly, Nicholas J. Kontos, George Kontos, Breanna J. Mueller, Samuel C. Norton, Joshua S. Godwin, Brad J. Schoenfeld, Melissa D. Boersma, Andrew D. Frugé, C. Brooks Mobley, Andreas N. Kavazis, Michael D. Roberts

**Author notes:** Address correspondence to: Michael D. Roberts, PhD, Professor & Director, Nutrabolt Applied and Molecular Physiology Laboratory School of Kinesiology, Auburn University, 301 Wire Road, Office 286, Auburn, AL 36849.

## Abstract

Skeletal muscle mass and training adaptations decline with aging, yet the proteomic basis of these attenuated responses remains unclear. We hypothesized that aging is accompanied by diminished proteome plasticity in response to resistance training (RT). The soluble proteome of VL biopsies was profiled in 17 younger (21.9 ± 2.5 yr) and 15 older (57.5 ± 6.9 yr) untrained males before and after 10–12 weeks of supervised RT using data-independent acquisition mass spectrometry (2,113 quantified proteins). At baseline, we detected 196 differentially expressed proteins (DEPs) significantly differed between age groups by Π-score (278 by FDR). A 5.6-fold difference in training-responsive was observed in younger vs. older adults (100 vs. 18 Π-score DEPs; 134 vs. 0 FDR-significant). Despite this quantitative attenuation, 61.6% of proteins changed in the same direction in both age groups (Spearman ρ = 0.284, p = 3.46 × 10⁻⁴⁰), indicating conserved but amplitude-compressed training responses (median |log₂FC|: 0.13 young vs. 0.09 old). RT in older adults partially reversed the aging proteome in that directionally different changes were observed in 75.2% of aging- or training-significant proteins in aging and training contrasts, with ribosomal and translational machinery showing the strongest reversal (cytoplasmic translation NES: −2.90 with aging, +2.60 with training). Ten WGCNA co-expression modules were identified, with age emerging as the dominant organizing principle (Turquoise module r-equiv = +0.59, p < 0.001). Module eigengenes discriminated age groups at the univariate level (Turquoise/Lipid Catabolism AUC = 0.96, q < 0.012), and training-induced module changes correlated with hypertrophic outcomes. Aging markedly attenuates but does not qualitatively alter skeletal muscle proteome plasticity. RT partially reverses aging proteome signatures, with translational machinery being the most responsive and mitochondrial programs the least responsive. Baseline proteomic state constrains adaptive capacity, suggesting that the molecular features distinguishing aging muscle directly may limit its hypertrophic response to RT.

## INTRODUCTION

Maintaining or increasing skeletal muscle mass is imperative among middle-aged and older adults for reducing the risk of life-threatening injuries from falls, cardiometabolic disease, and enhancing quality of life (1–3). Skeletal muscle mass typically peaks in the third to fourth decade and declines progressively thereafter, with losses accelerating after age 60 (4). This age-related loss (i.e., sarcopenia) affects up to 45% of adults over 70 years, representing a major public health concern with attributed healthcare costs of $40.4 billion annually (5, 6). Although resistance training (RT) is a viable strategy to maintain or increase muscle mass and strength, the hypertrophic responses are attenuated in older adults. This is evident in diminished changes in lean body mass relative to younger adults after equivalent training (7, 8) compounded by selective type II fiber atrophy that accounts for the majority of age-related muscle loss (9). This dissociation between preserved strength, achieved partly through enhanced neural adaptations (i.e., voluntary activation and motor unit recruitment) (10, 11), and attenuated hypertrophy (i.e., anabolic resistance) represents a fundamental barrier to exercise-based interventions for sarcopenia prevention (6).

At the molecular level, several mechanisms contribute to the attenuated hypertrophic response in older adults, many of which reflect a broader loss of proteostasis, a primary hallmark of biological aging (12). Aging may blunt activation of the mTORC1 pathway in response to mechanical loading and amino acid availability, reducing muscle protein synthetic rates (13), though the magnitude and consistency of these deficits remain debated (14, 15). Age-related deficits in autophagy-lysosomal and ubiquitin–proteasome systems impair protein quality control and promote accumulation of damaged proteins (16). Age-related deficits in mitochondrial oxidative capacity may also constrain bioenergetic resources needed for hypertrophic remodeling (17). Satellite cell dysfunction, characterized by a type II fiber–specific decline in satellite cell content and impaired activation in response to loading (18, 19), may compromise regenerative capacity. Similarly, reported deficits in cumulative protein synthesis coupled with attenuated ribosomal biogenesis may contribute to impaired RT responses in older individuals (20).

We put forth a recent hypothesis to suggest that aging is associated with reduced “proteome plasticity” to RT (21); this being a term we use herein to denote the capacity of the muscle proteome to dynamically respond to anabolic stimuli, operationalized as the number, magnitude, and functional coherence of training-induced protein abundance changes. Several lines of evidence support this concept. For instance, baseline proteomic differences between younger and older muscle are substantial (21–27), including mitochondrial metabolism, molecular chaperones, and cytoskeletal elements. These baseline differences could factor into the diminished training adaptations and constrained plasticity of the aged proteome. Chronic transcriptomic responses to RT are similarly dampened in older adults, with seven-fold fewer responsive transcripts in older type II fibers (28), Nevertheless, RT can partially reverse age-related gene expression signatures (29). Pre-training transcriptomic profiles predict response heterogeneity (30), and ribosomal production predicts hypertrophy in older adults (31), suggesting that aging constrains both transcriptional and translational plasticity. However, direct comparisons of training-induced proteome remodeling between younger and older adults remain scarce.

Among the limited studies examining RT-induced proteomic changes across age groups, limitations include designs that did not train both age groups for direct comparison (21), an emphasis on muscle function (rather than size changes) and proteomic outcomes (32), or limited dynamic range detection with first generation proteomic interrogation techniques (22). Deep proteomics studies of exercise-induced skeletal muscle adaptations have arisen recently with notable contributions from the MoTrPAC consortium (33), and independent research laboratories including our own (21, 26, 32, 34–36).

The present investigation was motivated by the sparse evidence indicating that RT-induced proteome remodeling may differ between younger and older adults. Herein, the soluble proteome of vastus lateralis (VL) biopsies was profiled in 17 younger (21.9 ± 2.5 yr) and 15 older (57.5 ± 6.9 yr) untrained males before and after 10–12 weeks of supervised RT using data-independent acquisition mass spectrometry (2,113 quantified proteins). We hypothesized that skeletal muscle proteome plasticity in response to RT would be constrained in the older versus younger cohort. We further hypothesized that age-related constraints would extend to the pathway level, manifesting as discordance in training adaptations between age groups. Drawing on evidence that RT can partially reverse aging gene expression signatures, we hypothesized that RT might similarly reverse age-related proteome signatures, suggesting that at least some age-related alterations represent modifiable molecular phenotypes rather than irreversible features of aging. Beyond differential expression, we sought to identify trajectory clusters of co-regulated proteins and to determine whether baseline proteomic profiles predict training-induced hypertrophic outcomes.

## METHODS

### Participants and ethical approval

This study is a secondary analysis of VL muscle biopsies obtained from three parent investigations conducted at Auburn University (37–39). All parent studies were approved by the Auburn University Institutional Review Board (Protocol numbers 19-249, 20-136, & 24-863) and conducted in accordance with the Declaration of Helsinki except for two studies (20-136 & 24-863) not being registered as clinical trials. All participants provided written informed consent prior to enrollment.

A total of 17 younger (21.9 ± 2.5 yr, range: 19–26 yr) and 15 older (57.5 ± 6.9 yr, range: 42–69 yr) untrained males were included. All participants had refrained from structured resistance training for ≥6 months prior to enrollment, had a body mass index between 18.5 and 35 kg/m², had no contraindications to muscle biopsy or resistance exercise, and were not using medications known to affect muscle metabolism (e.g., corticosteroids, testosterone, growth hormone). Two individual timepoint samples from separate younger participants were flagged and removed by a four-method consensus (per-sample missingness, PCA Mahalanobis distance, median-intensity MAD, and inter-sample Pearson correlation), with the unpaired partner from each participant retained, yielding 62 analyzed samples from 17 younger and 15 older adults. Participant characteristics are presented in Table 1, and excluded samples are identified in S1 Table. As a secondary analysis, sample sizes were determined by the parent investigations. Post hoc analysis (pwr.t.test; 80% power, α = 0.10) indicated that the paired within-group contrasts could detect effects of d ≥ 0.65 (log₂FC ≥ 0.37), while the unpaired Aging and Interaction contrasts required d ≥ 0.93 (log₂FC ≥ 0.63) (S6 Table).

**Table 1.** Pre-intervention characteristics.

| Variable | Younger (mean ± SD) | Older (mean ± SD) | p-value |
| --- | --- | --- | --- |
| Age (years) | 21.9 ± 2.5 | 57.5 ± 6.9 | <0.001 |
| Body mass index (kg/m<sup>2</sup>) | 24.5 ± 2.8 | 28.7 ± 2.9 | <0.001 |
| DXA LBM (kg) | 56.6 ± 5.7 | 58.3 ± 6.5 | 0.431 |
| VL thickness (cm) | 2.08 ± 0.60 | 1.95 ± 0.41 | 0.459 |

Abbreviations: DXA, dual-energy x-ray absorptiometry; LBM, lean body mass (bone-free); VL, vastus lateralis

### Training protocol

All participants completed supervised, progressive resistance training programs targeting major muscle groups. Training was conducted twice per week over 10–12 weeks under direct supervision by trained research personnel. Specific training durations and programming details have been described in the respective parent investigations. Briefly, sessions included lower-body exercises (45° leg press, leg extensions, lying leg curls, and squat or hex-bar deadlift variations) and upper-body exercises (chest press variations and cable pull-downs or rows).

Participants performed 3–5 sets of 6–15 repetitions per exercise with progressive overload guided by ratings of perceived exertion or repetitions in reserve. Volume load (sets × repetitions × load) was tracked throughout the intervention and compared between age groups. Volume load, lean body mass, and VL thickness were available for all participants (n = 17 younger, n = 15 older).

### Vastus Lateralis Muscle Thickness

VL muscle thickness was assessed using B-mode ultrasound (NextGen LOGIQe R8: GE Healthcare) with a multifrequency linear-array transducer (L4-12T, 4 – 12 MHz, GE Healthcare). Briefly, participants laid supine on an athletic training table while a technician oriented the probe in the transverse plane on the lateral aspect of their thigh to obtain still images of the VL. The pre-intervention biopsy scar was used as a guide for replicating pre- and post-intervention measurements. Ultrasound settings were held constant across participants for each individual study. VL muscle thickness was quantified by measuring the linear distance from the deep aponeurosis to the superficial interface between the VL and the subcutaneous adipose tissue in ImageJ (National Institutes of Health, Bethesda, MD).

### Body Composition Assessment

The height and weight of all participants was assessed using a digital column scale Seca 769; Hanover, MD). Prior to body composition testing, participants submitted a urine sample (≥5 mL) which was assessed for urine specific gravity using a handheld refractometer (ATAGO: Bellevue, WA). Across studies, urine specific gravity was ≤1.030 for all participants analyzed. Participants then laid supine for an assessment of body composition via dual-energy X-ray absorptiometry (DXA) (Lunar Prodigy; GE Corporation; Fairfield, CT) from which bone-free lean body mass (LBM) was derived.

### Tissue collection and processing

VL muscle biopsies were obtained at baseline (PRE) and ≥72 hours following the final training session (POST) using the 5-gauge percutaneous needle biopsy technique with suction. The mid-belly of the right VL was identified at the midpoint between the inguinal crease and proximal border of the patella, locally anesthetized with 1% lidocaine, and approximately 50–100 mg of tissue was collected. Immediately following collection, muscle samples were cleared of visible blood, adipose, and connective tissue, and snap-frozen in liquid nitrogen. Samples were stored at−80°C until processing. For proteomic analysis, the soluble protein fraction was extracted using a 1X general tissue lysis buffer (Cell Signaling Technology; Danvers, MA, USA; catalog #: 9803) containing 20 mM Tris-HCl (pH 7.5), 150 mM NaCl, 1 mM Na2EDTA, 1 mM EGTA, 1%

Triton, 2.5 mM sodium pyrophosphate, 1 mM beta-glycerophosphate, 1 mM Na3VO4, 1 µg/ml leupeptin. Briefly, ∼20 mg muscle was minced on a liquid nitrogen-cooled mortar and placed in 1.7 mL tubes on ice containing 500 µL of buffer. Tissue was mechanically homogenized with tight-fitting pestles, and samples were centrifuged at 500 × g for 5 minutes. Following centrifugation, supernatants (termed ‘lysates’ hereafter) were placed in new tubes and frozen at −80°C until peptide generation. Notably, this lysis procedure tends to solubilize membrane, cytosolic, nuclear, and some contractile and extracellular matrix proteins according to prior assessments from our laboratory (40, 41), thus yielding a broad-range spectrum of proteins across all subcellular compartments.

### Proteomic analysis

Protein lysates (50 µg per sample) were prepared for LC-MS/MS analysis using EasyPep Mini MS Sample Prep Kit (Thermo Fisher; Waltham, MA, USA; catalog #: A4006). Briefly, samples were transferred into new 1.7 mL microcentrifuge tubes and final volumes were adjusted to 100 µL with general cell lysis buffer (Cell Signaling). Reduction (50 µL) and alkylation (50 µL) solutions were added to samples, gently mixed, and incubated at 95°C using a heat block for 10 minutes. After incubation, the sample was removed from the heat block and cooled to room temperature. A reconstituted enzyme Trypsin/Lys-C Protease solution (25 µL) was added to the reduced and alkylated protein sample and incubated with shaking at 37°C for 2 hours to digest proteins. Digestion stop solution (25 µL) was then added to samples and peptides were cleaned using a peptide clean-up column according to the kit instructions.

An externally calibrated Thermo Orbitrap Exploris 240 (high-resolution electrospray tandem mass spectrometer) was used in conjunction with Dionex UltiMate3000 RSLC Nano System (Thermo Fisher) for proteomics. Samples were aspirated into a 20 µL loop and loaded onto the trap column (Acclaim PepMap 100 nanoviper tubing 75 µm i.d. × 2 cm; Thermo Fisher). The flow rate was set to 300 nL/min for separation on the analytical column (Easy-Spray pepmap RSLC C18 50 µM × 15 cm; Thermo Fisher). Mobile phase A was composed of 99.9% H2O (EMD Omni Solvent; Millipore, Austin, TX, USA), and 0.1% formic acid and mobile phase B was composed of 80% acetonitrile and 0.1% formic acid. A 135-minute linear gradient from 3% to 50% B was performed. The eluent was directly nanosprayed into the mass-spectrometer. During chromatographic separation, the Orbitrap Exploris was operated in a data-independent mode and under direct control of the Thermo Xcalibur 4.4.16.14 (Thermo Fisher). The MS data were acquired from (380 to 985 m/z) at 60,000 resolution. MS2 spectra were acquired in data-independent acquisition mode with a 10 m/z isolation window, 1 m/z overlap, collision energy of 28, and a resolution of 15,000. All measurements were performed at room temperature, and three technical replicates were run for each sample. Raw .dia files acquired from mass spectrometry were processed using DIA-NN (v 2.2.0) (42). All raw files were analyzed jointly in a single batch to enable retention-time alignment and match-between-runs (MBR) quantification. In silico digestion was performed with trypsin-P specificity, allowing one missed cleavage. Carbamidomethylation of cysteine was specified as a fixed modification, while oxidation of methionine was included as a variable modification, with one variable modification permitted per peptide. N-terminal methionine excision was enabled. Peptide identification was guided by the DPHLv2 spectral library (43). For protein reannotation, a reviewed *Homo sapiens* UniProt reference proteome (release 2020-04) supplemented with indexed retention-time (iRT) calibration peptides, reversed decoys, and common contaminants were used. Peptide and protein-level false discovery rates were controlled at 1%. Data-independent acquisition (DIA) was chosen for its superior quantitative reproducibility and depth in skeletal muscle proteomics (44).

Proteins annotated as expressed in skeletal muscle (Human Protein Atlas) were deduplicated by UniProt accession and retained if detected in ≥10 samples within at least one experimental group. Highly abundant blood proteins and common contaminants were removed. Sample outliers were identified by a four-method consensus (per-sample missingness, PCA Mahalanobis distance, median-intensity MAD, and inter-sample Pearson correlation); two individual timepoint samples were removed and the unpaired partners retained (excluded samples identified in S1 Table). Abundances were normalized by cyclic loess via the proteoDA R package (Thurman et al., https://github.com/ByrumLab/proteoDA). After filtering, 2,113 proteins were retained for downstream analysis. The quality-control pipeline is summarized in S1 Figure. Raw (S2 Table), normalized (S3 Table), and imputed (S4 Table) protein matrices are provided as supplementary data.

### Statistical analysis

Differentially expressed protein (DEP) abundance was assessed using the limma R package with empirical Bayes moderation (45, 46). The model accounted for the paired PRE–POST design using the duplicateCorrelation function to estimate within-subject correlation. Four contrasts were defined: Training in Young (Young_Post − Young_Pre), Training in Old (Old_Post − Old_Pre), Aging at Baseline (Old_Pre − Young_Pre), and Age × Training Interaction ([Old_Post − Old_Pre] − [Young_Post − Young_Pre]). Significance was determined using the Π-score, which is a composite metric integrating both statistical significance and fold-change magnitude (Π = p^|log₂FC|^), with proteins considered significant at Π < 0.05. Significance was also assessed by Benjamini–Hochberg-corrected false discovery rate (FDR < 0.05) and nominal p-value (p < 0.05) to provide complementary thresholds. Missing values were imputed with missForest (ntree = 100, maxiter = 10) (47), selected 17-method benchmark that scored candidate methods against the non-imputed reference on fold-change preservation, pathway-level concordance, and reconstruction error; missforest was chosen for its strong composite performance, established literature precedent, and straightforward implementation (S5 Table). Differential abundance testing was performed on non-imputed data, as limma fits each protein independently using only available observations. Imputed values were employed where a complete data matrix was required (PCA, WGCNA). Additionally, the fry gene-set rotation test (limma::fry) was used to assess whether pre-defined DEP sets showed concordant or reversed behavior in an independent contrast, accounting for inter-gene correlation via the duplicateCorrelation estimate. Module-trait associations from weighted gene co-expression network analysis (WGCNA) were assessed via linear mixed models (eigengene ∼ group + (1|subject)) with Kenward-Roger degrees of freedom to account for the paired design (48, 49), and effect sizes reported as equivalent correlations derived from the model t-statistics.

#### Protein set enrichment analysis (PSEA)

Two primary approaches were used to convey pathway-level information. First, ranked-list enrichment (fgseaMultilevel) tested proteins ordered by the limma moderated t-statistic against MSigDB Hallmark, KEGG Medicus, Reactome, GO:BP, and GO Slim gene sets (size 15–500; disease/cancer terms excluded), with Benjamini–Hochberg correction applied per database (50). Second, concordance and reversal analyses compared contrast-level NES values across an a priori Hallmark and GO Slim collection. Over-representation analysis (fgsea::fora) was applied to proteins partitioned by scatter-plot quadrant with Benjamini–Hochberg correction and Jaccard deduplication (cutoff = 0.5) to reduce redundancy among overlapping gene sets (51).

#### Concordance analysis

To assess whether younger and older adults exhibit concordant proteomic responses to training, a Spearman rank correlation was computed between Training Effect logFC values across all quantified proteins. Proteins were further classified into four concordance categories based on the significance and direction of their training response in each age group. Threshold-free concordance was assessed using Rank-Rank Hypergeometric Overlap (RRHO2), which partitions proteins into quadrants of concordant and discordant regulation without reliance on significance cutoffs (51). Global proteomic composition was visualized by principal component analysis on the imputed matrix, and the contribution of age, training time point, and their interaction was quantified by permutational multivariate analysis of variance (PERMANOVA; adonis2, 999 permutations) on Euclidean distances (52). Per-contrast effect-size distributions were compared using pairwise Wilcoxon rank-sum tests on absolute log2 fold-change (log₂FC) values.

#### Reversal analysis

To evaluate whether resistance training reverses age-related proteomic changes, each protein’s aging effect (Old_Pre − Young_Pre) was compared with its training response in older adults (Old_Post − Old_Pre). Proteins were classified as reversed (training-induced change opposes aging effect), exacerbated (training reinforces aging effect), or negligible (|log₂FC| < 0.2) based on the direction and magnitude of these opposing effects. DEPs were further classified into four directional categories based on whether the training response in older adults opposed or reinforced the aging effect.

#### Software and data availability

All statistical analyses and visualizations were performed in R (version 4.5.2). Key packages included limma, proteoDA, missForest, fgsea, msigdbr, GO.db, WGCNA, ggplot2, and patchwork. Code and processed data are available at https://github.com/Dustyn-T-Lewis/YvO_2025. Source data for all main figures are provided in S7–S13 Tables.

## RESULTS

### Participant characteristics and training outcomes

Volume load was similar between age groups (p = 0.771; Fig. 1A). Both age groups increased DXA-derived lean body mass following training (Young: +2.21 kg, Older: +1.18 kg; between-group p = 0.179; Fig. 1B) and VL thickness (Younger: +0.47 cm, Old: +0.20 cm; Fig. 1C). Young adults gained significantly more VL thickness than older adults (Δ = 0.263 cm, p = 0.003), confirming differential hypertrophic response despite comparable training stimulus. Younger adults also showed greater gains in deadlift one-repetition maximum (interaction p < 0.001; n = 17 young, 9 old) and Type II fiber cross-sectional area (interaction p = 0.01; n = 13 young, 10 old; S2 Figure).

**Figure 1.**
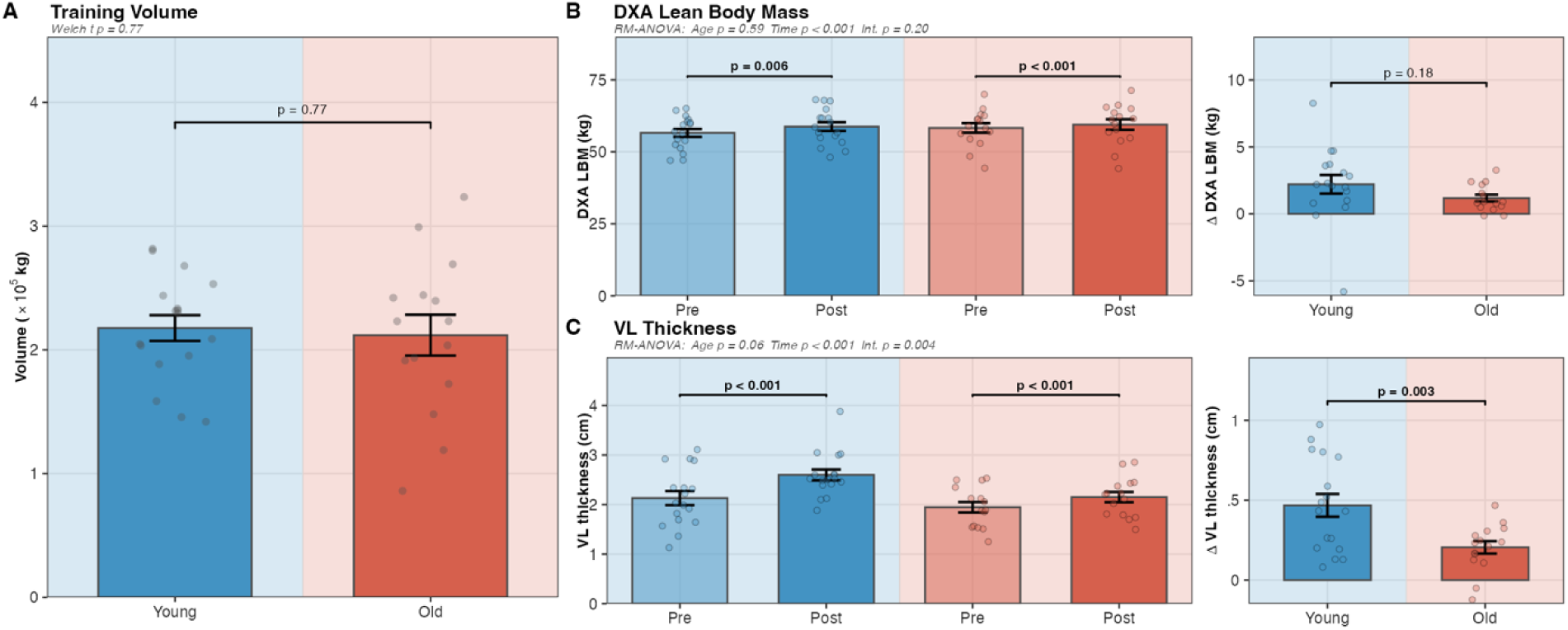
Phenotypic responses to resistance training in younger (n = 17) and older (n = 15) adults. (A) Volume load; groups were similar (p = 0.771). (B) DXA lean body mass pre- and post-training (left) and between-group change (right); both groups gained LBM (Young p = 0.006; Old p < 0.001); between-group Δ was not significant (p = 0.179). RM-ANOVA: Age p = 0.592, Time p < 0.001, Interaction p = 0.196. (C) VL thickness pre- and post-training (left) and between-group change (right); younger adults gained more than older adults (Δ = 0.263 cm, p = 0.003). RM-ANOVA: Age p = 0.060, Time p < 0.001, Interaction p = 0.004. Values are means ± SE. DXA, dual-energy X-ray absorptiometry; LBM, lean body mass; VL, vastus lateralis.

### Proteomics summary

After quality-control filtering, 2,113 proteins were quantified across 62 VL biopsies from 17 younger and 15 older adults sampled before and after resistance training. Median inter-individual coefficients of variation ranged from 23–26% across all four groups (S3 Figure). Baseline CVs were comparable between age groups, though training induced modest but significant shifts in variability, with the largest divergence occurring post-training. Principal component analysis revealed partial separation by age group, though not by training status (Fig. 2A). PERMANOVA nonetheless detected significant main effects of age (*R*^2^ = 0.089, p = 0.001) and training time point (*R*^2^ = 0.029, p = 0.001), while the age × time interaction was non-significant (*R*^2^ = 0.013, p = 0.282). Effect-size distributions differed substantially across contrasts (all pairwise p < 0.001, Wilcoxon rank-sum on | log₂FC|; Fig. 2B). The Aging contrast exhibited the broadest spread (median |log₂FC| = 0.18), followed by Training in Young (0.13) and Training in Old (0.09), revealing progressive compression of per-protein effect sizes from the aging signature to the training response in older adults.

**Figure 2.**
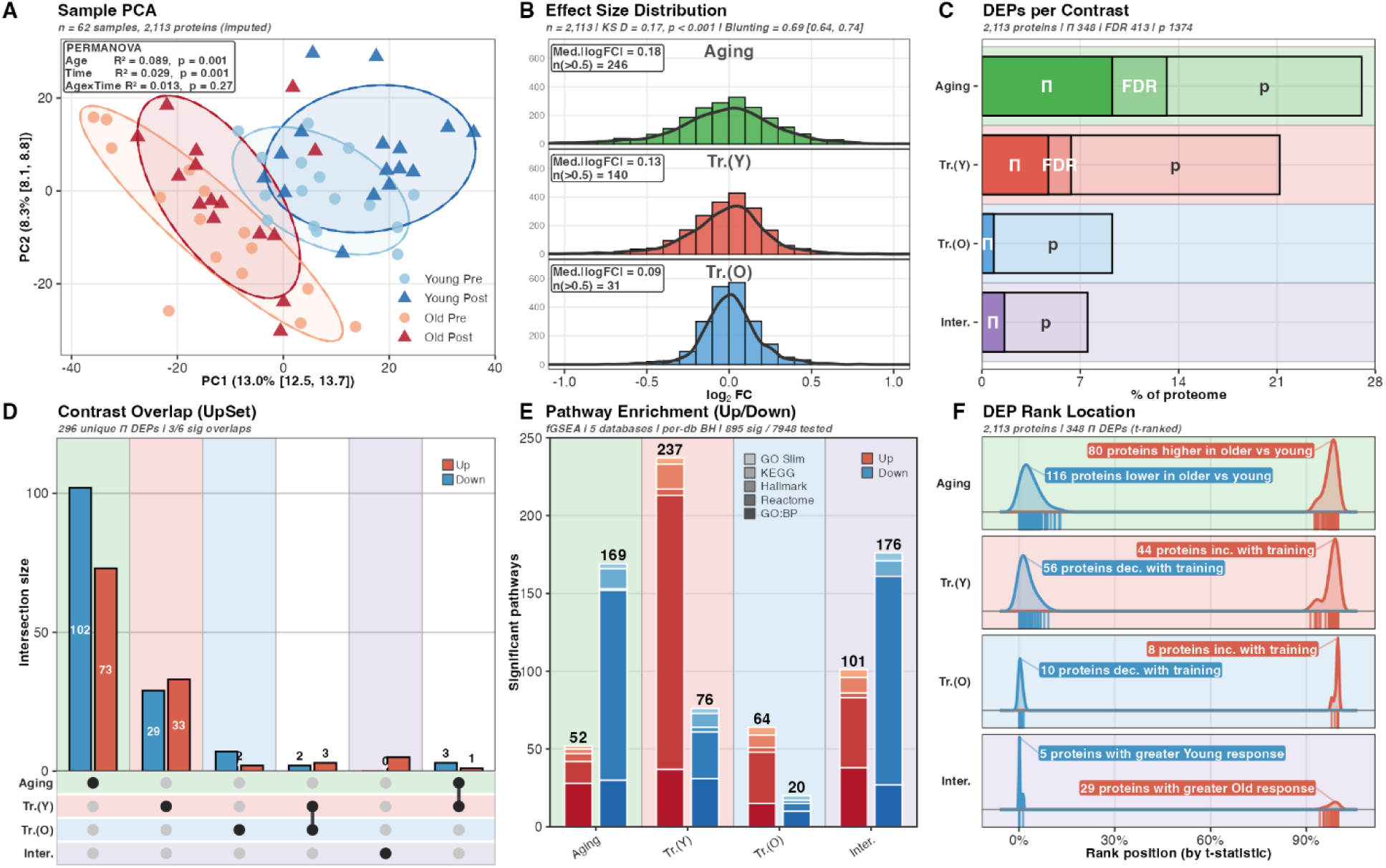
Proteome overview and differential expression landscape (2,113 proteins, 62 samples). (A) PCA of the imputed protein matrix; PERMANOVA: Age R² = 0.089 (p = 0.001), Time R² = 0.029 (p = 0.001), Age×Time R² = 0.013 (p = 0.282). (B) Per-contrast |log₂FC| distributions showing progressive compression from Aging (median 0.18) to Training in Young (0.13) to Training in Old (0.09). (C) DEP counts at three significance thresholds (Π-score, FDR, nominal p). (D) UpSet intersection of Π-score DEPs across contrasts. (E) Fraction of fGSEA-enriched pathways (BH-corrected) across Hallmark, GO Slim, GO:BP, KEGG, and Reactome gene sets, separated by enrichment direction. (F) DEP rank location within the t-statistic–ranked proteome per contrast. DEP, differentially expressed protein; FC, fold change; PCA, principal component analysis.

DEP analysis at three significance thresholds revealed stark contrast-dependent asymmetry (Fig. 2C; diagnostics in S4 Figure, per-protein results in S6 Table). The Aging contrast yielded the most DEPs at every threshold (196 at Π < 0.05; 278 at FDR < 0.05; 571 at p < 0.05), followed by Training in Young (100; 134; 448). Training in Old produced strikingly few DEPs (18 at Π < 0.05; 0 at FDR < 0.05)—a 5.6-fold reduction relative to younger adults. The interaction contrast identified 34 DEPs (Π < 0.05), confirming that the attenuated older response is statistically robust. Among the 196 aging-significant proteins, the most strongly depleted in older muscle were sarcomeric components (LMOD2, log₂FC = −2.35; ACTN3, −2.12), while the most elevated included GPT2 (+1.68, amino acid catabolism) and PLIN2 (+1.12, lipid droplet storage). Full per-protein results for all contrasts are provided in S6 Table.

UpSet intersection analysis of the 296 unique Π-score DEPs revealed predominantly contrast-specific signatures (Fig. 2D), with Aging contributing the largest exclusive set (175 proteins, 59%), followed by Training in Young (62, 21%), and minimal higher-order intersections. Protein set enrichment analysis (fGSEA; Hallmark, GO Slim, GO:BP, KEGG, and Reactome gene sets with BH correction per database) identified 895 significantly enriched pathway-contrast combinations across all four contrasts (padj < 0.05; Fig. 2E). The rank-position distribution of Π-score DEPs within the t-statistic–ordered proteome confirmed that significant proteins cluster at the extremes of each contrast (Fig. 2F). The Training Young and Interaction contrasts yielded the most pathway-level signal (313 and 277 pathways, respectively), while Training in Old yielded 3.7-fold fewer enriched pathways (84 vs. 313). Notably, the Interaction contrast—despite only 34 individual DEPs—identified substantial pathway-level signal (277 pathways), indicating that age-dependent training differences manifest as coordinated shifts across functionally related proteins.

### Training responses are directionally conserved but magnitude-compressed in older adults

To assess whether younger and older adults mount qualitatively similar or distinct training responses, we compared protein-level changes between age groups. Flanked volcano plots revealed 100 Π-score DEPs in younger versus 18 in older adults (Fig. 3B–C). Despite this quantitative disparity, pathway-level insets showed overlapping enrichment themes in both age groups, though younger adults exhibited pronounced oxidative phosphorylation suppression absent in older adults.

**Figure 3.**
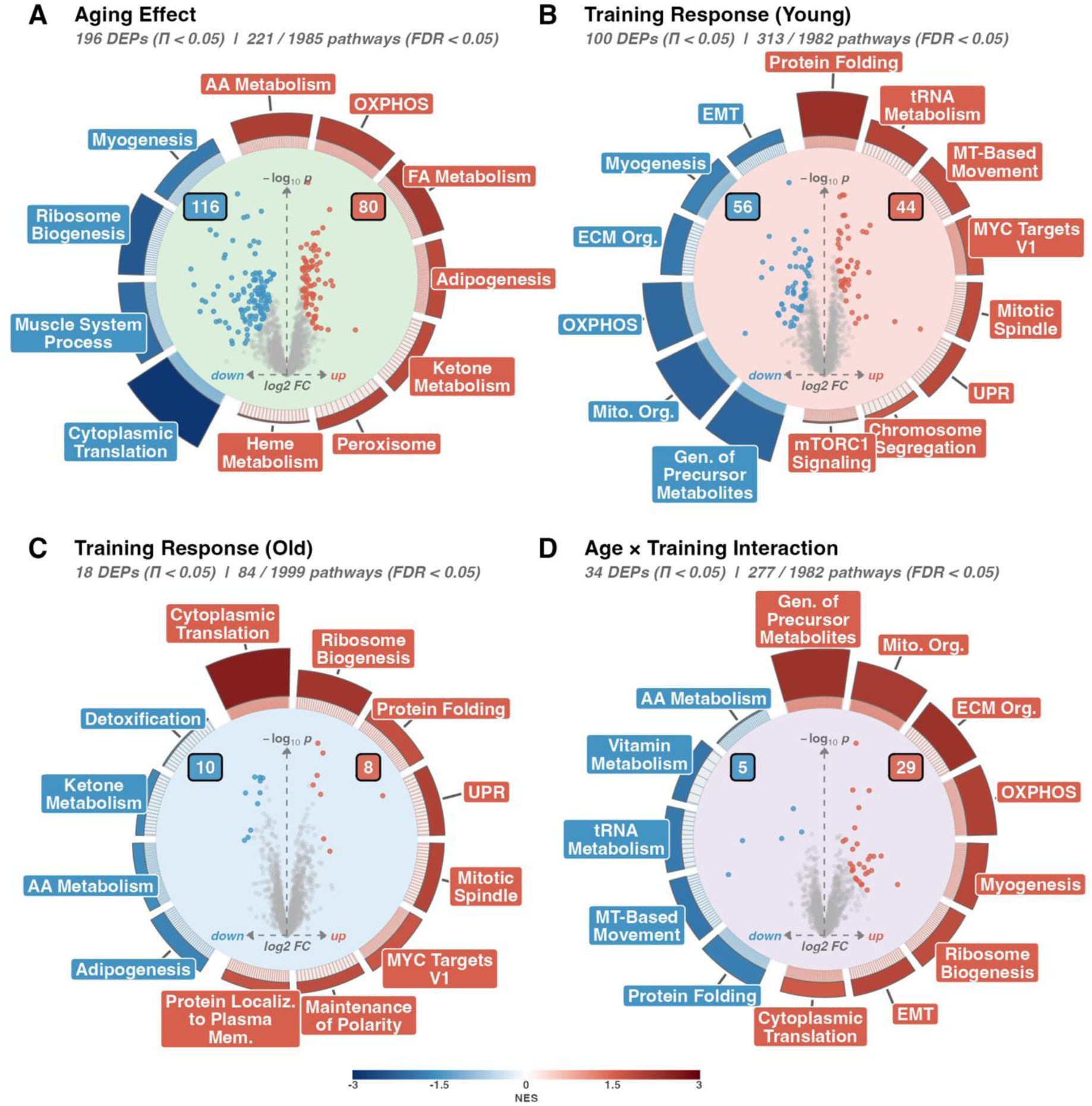
Volcano ring plots for the four primary contrasts. (A) Aging effect (Old_Pre − Young_Pre; 196 Π-score DEPs; 221 enriched pathways). (B) Training in Young (Young_Post − Young_Pre; 100 DEPs; 313 pathways). (C) Training in Old (Old_Post − Old_Pre; 18 DEPs; 84 pathways). (D) Age × Training interaction ([Old_Post − Old_Pre] − [Young_Post − Young_Pre]; 34 DEPs; 277 pathways). Inner scatter plots show protein-level log₂FC vs. −log₁₀(p); red, upregulated; blue, downregulated. Outer ring insets display the top fGSEA-enriched pathways coloured by normalised enrichment score (NES). DEP significance threshold: Π-score < 0.05. DEP, differentially expressed protein; NES, normalised enrichment score.

Training-induced log₂FC values showed a weak but highly significant positive correlation between age groups (Spearman ρ = 0.284, p = 3.46 × 10^-40^; Fig. 4A), with 61.6% of proteins changing in the same direction. All six proteins reaching Π-score significance in both age groups were directionally concordant. Effect sizes were systematically compressed in older adults (median |log₂FC|: 0.13 vs. 0.09), consistent with an intact but attenuated training response. A protein-permuted null confirmed this correlation was not circular (null mean ≈ 0, p < 0.001), and reversal classification was stable across |log₂FC| thresholds (S8 Figure).

**Figure 4.**
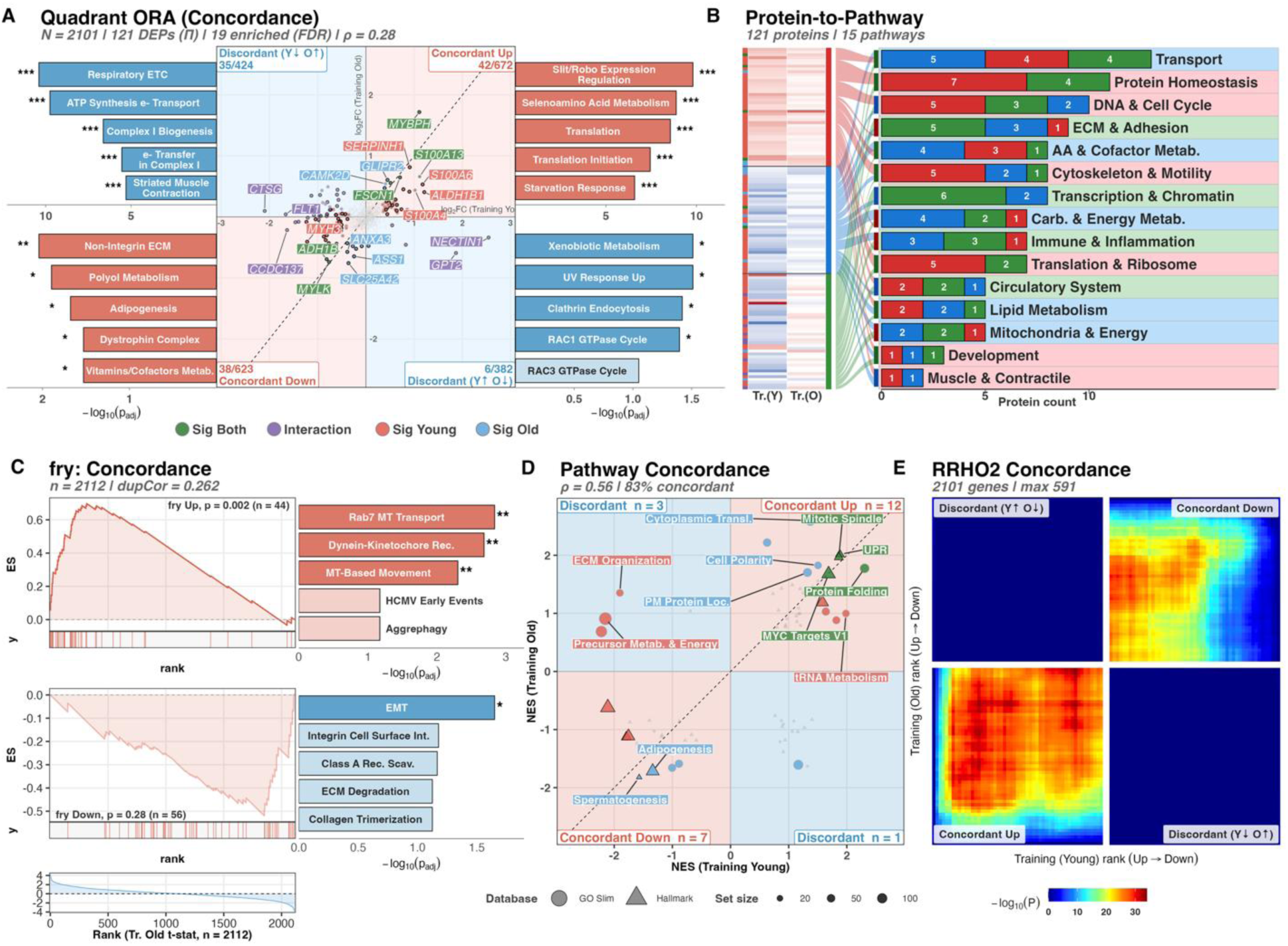
Training response concordance between younger and older adults. (A) Quadrant scatter of Training-in-Young vs. Training-in-Old log₂FC (2,101 proteins); 121 met Π < 0.05; 19 ORA-enriched pathways at FDR < 0.05; Spearman ρ = 0.284 (p = 3.46 × 10⁻⁴⁰). (B) Response-pattern heatmap of 121 significant proteins across 15 GO Slim categories, partitioned into concordant and discordant classes. (C) fry gene-set rotation test of Young Π-DEPs against the Training-in-Old t-statistic; Young-up p = 0.002 (n = 44), Young-down p = 0.28 (n = 56). (D) Pathway-level NES concordance across 69 a priori Hallmark/GO Slim sets; Spearman ρ = 0.564 (p = 4.40 × 10⁻⁷); 23 pathways significant. (E) Threshold-free RRHO2 concordance map. FC, fold change; NES, normalised enrichment score; ORA, over-representation analysis.

RRHO2 analysis confirmed this pattern without reliance on arbitrary significance thresholds (Fig. 4E), with strong concordance signals in both upregulated and downregulated quadrants and negligible discordant signal, indicating that the dominant pattern is concordance between age groups.

Pathway-level concordance was stronger than protein-level concordance (Spearman ρ = 0.56, p = 4.40 × 10⁻⁷; Fig. 4D). Four pathways reached significance in both age groups, all directionally concordant. The largest age-divergent signal was in oxidative phosphorylation (Hallmark; NES: −2.11 in young vs. −0.62 in old), with 23 of 69 curated Hallmark and GO Slim pathways reaching significance in at least one age group (concordance diagnostics in S8 Figure).

Among the 121 proteins reaching Π-score significance in any training or interaction contrast (Fig. 4A), 42 showed concordant upregulation, 38 concordant downregulation, and 41 discordant responses between age groups. Concordantly downregulated proteins were enriched for extracellular matrix organization (padj < 0.001) and collagen biosynthesis, suggesting that ECM remodeling represents a shared prerequisite for training-induced fiber adaptation regardless of age. A rotation gene set test (fry) confirmed that proteins upregulated by training in younger adults (n = 44 Π-DEPs) responded concordantly in older adults (p = 0.002; Fig. 4C), consistent with conservation of training direction despite magnitude compression.

### Resistance training partially reverses the aging proteome signature

Having established that training responses are directionally concordant but magnitude-compressed age groups, we next examined whether resistance training reverses age-related proteomic changes. Comparing the Aging contrast (196 Π-score DEPs; Fig. 3A) with the Training-in-Old contrast revealed 158 of 210 Π-score DEPs with opposite-sign changes between the Aging and Training-in-Old contrasts (75.2%) (Fig. 3C). Pathway-level ring insets showed a striking mirror image in that pathways downregulated with aging were upregulated with training, and vice versa. Of the 210 proteins reaching Π-score significance across the Aging or Training-in-Old contrasts, 158 (75.2%) fell in reversal quadrants (Fig. 5A), indicating that training in older adults preferentially targets aging-dysregulated proteins. RRHO2 analysis revealed strong enrichment in both reversal quadrants of the reversal map (Fig. 5E). The strongest signal occurred in the Aging-Down/Training-Up quadrant (max −log10 p = 35.9), with reciprocal reversal also significant (−log10 p = 29.0). Exacerbation signals were negligible, indicating that training selectively reversed rather than exacerbated age-related proteomic changes. A protein-permuted null confirmed this correlation was not circular (null mean ≈ 0, p < 0.001), and reversal classification was stable across |log₂FC| thresholds (S9 Figure).

**Figure 5.**
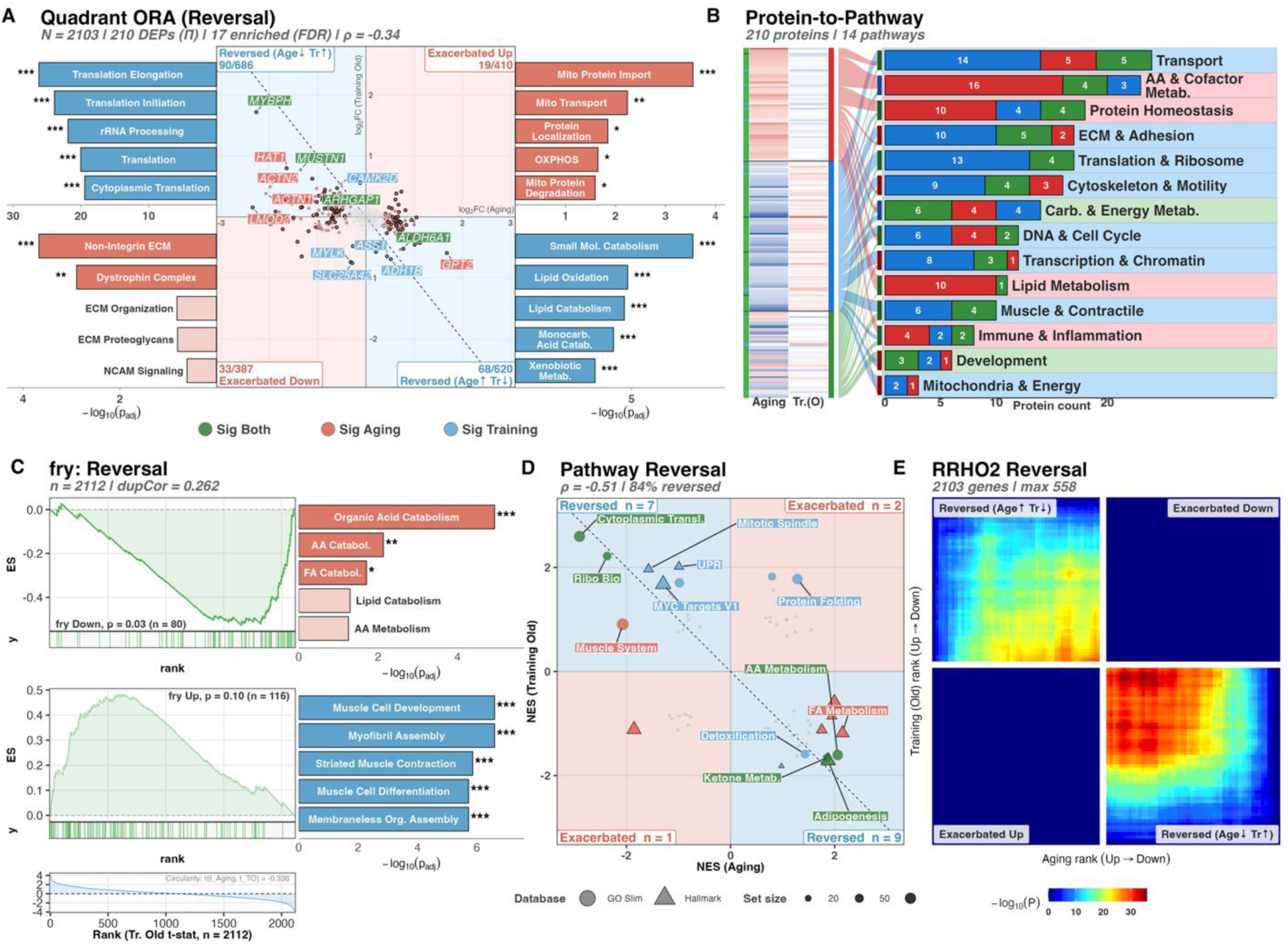
Resistance training in older adults partially reverses the aging proteome signature. (A) Quadrant scatter of Aging vs. Training-in-Old log₂FC (2,103 proteins); 210 met Π < 0.05; 17 ORA-enriched pathways at FDR < 0.05; Pearson r = −0.45 among 210 Π-significant proteins. (B) Reversal-pattern heatmap of 210 significant proteins across 14 GO Slim categories, partitioned into reversed and non-reversed classes. (C) fry gene-set rotation test: Aging-up proteins reversed toward lower abundance with training (p = 0.03, n = 80); Aging-down trended higher (p = 0.10, n = 116). (D) Pathway-level NES reversal across 69 a priori Hallmark/GO Slim sets; Spearman ρ = −0.51; 19 significant, 84% reversed. (E) Threshold-free RRHO2 reversal map. FC, fold change; NES, normalised enrichment score; ORA, over-representation analysis.

At the pathway level, 19 pathways showed significant enrichment in either or both contrasts (Fig. 5D), with cytoplasmic translation showing the strongest reversal (NES: −2.90 with aging, +2.60 with training). Among the 30 highest-magnitude aging DEPs (Fig. 5A–B), the majority showed directional reversal, enriched for muscle cell development and myogenesis, indicating that the structural proteins most affected by aging are precisely those most responsive to training-mediated recovery. Taken together, the reversal analysis indicates that training in older adults directionally opposes aging-related proteomic changes, with the strongest reversal in ribosomal and translational machinery and the weakest in oxidative phosphorylation and lipid catabolism pathways.

### Weighted co-expression network analysis reveals age- and training-associated protein modules

We constructed a signed WGCNA network from 2,113 proteins across 62 samples, identifying 10 co-expression modules (67–345 proteins; 1,606 assigned, 507 unassigned; Fig. 6A). Network construction used Pearson correlation with soft-thresholding power 12 (scale-free *R²* = 0.874; threshold relaxed to 0.85 from the conventional 0.90 per Langfelder & Horvath guidance for small-sample studies). Because paired pre/post samples violate the independence assumption of standard module-trait correlations, associations were assessed via linear mixed models (eigengene ∼ group + (1|subject)) with Kenward-Roger degrees of freedom. Age was the dominant organizing principle, with three modules reaching BH-corrected significance: Turquoise (345 proteins; lipid catabolism; r = +0.59, pBH = 4.6 × 10⁻⁵), Magenta (78; translation initiation; r = −0.53, pBH = 5.7 × 10⁻⁴), and Brown (197; muscle contraction; r = −0.44, pBH = 0.008). The Blue module (261; cell cycle/proteostasis) was the only module significantly associated with training in younger adults (r = +0.63, pBH = 0.002). No modules reached significance for Training in Old or the Interaction contrast. Among stratified within-group correlations, significant phenotypic association was only observed for the Green module (164; oxidative phosphorylation): eigengene change was inversely correlated with training-induced VL thickness change in younger adults (r = −0.69, pBH = 0.046; corrected across 11 modules within this stratum). Among the remaining stratified correlations (none BH-significant), the strongest per-trait associations were: Blue (cell cycle/proteostasis) with baseline body composition in younger adults (BMI r = −0.55, LBM r = −0.56), Red (immunity/heme) with baseline BMI in older adults (r = +0.55), and Brown (muscle contraction) with training-induced LBM change in younger adults (r = −0.62).

**Figure 6.**
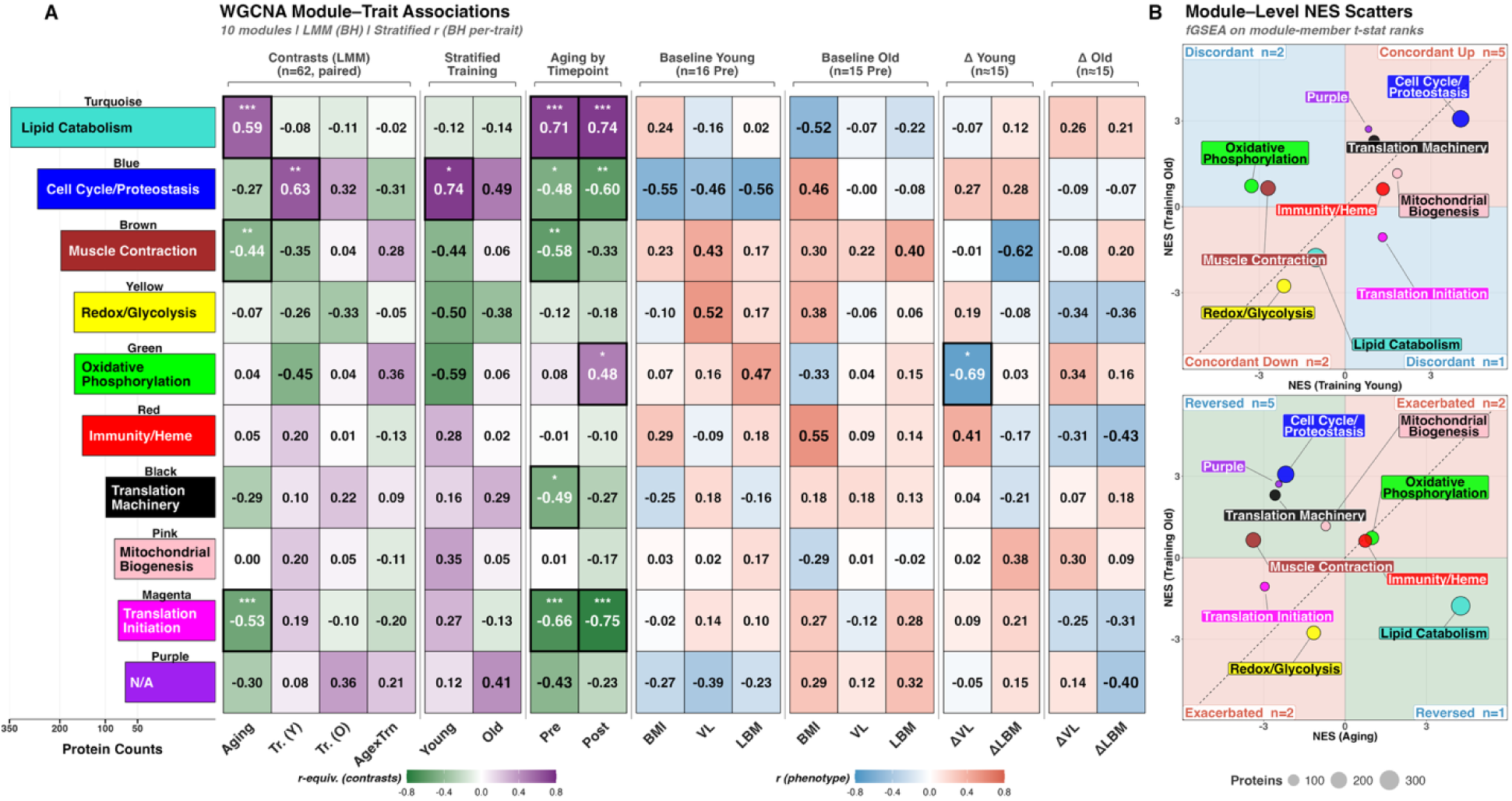
WGCNA co-expression network analysis of 2,113 proteins across 62 samples. (A) Module–trait association heatmap for ten modules. Left columns: r-equivalents from paired linear mixed models (Aging, Training in Young, Training in Old, Age×Training); right columns: stratified Pearson correlations with baseline phenotypes and training-induced changes per age group. BH-corrected significance and module sizes annotated. Three modules were age-associated: Turquoise/Lipid Catabolism (r = +0.59, p_BH = 4.6 × 10⁻⁵), Magenta/Translation Initiation (r = −0.53, p_BH = 5.7 × 10⁻⁴), Brown/Muscle Contraction (r = −0.44, p_BH = 0.008). (B) Module-level NES for Training-in-Young vs. Training-in-Old. (C) Module-level NES for Aging vs. Training-in-Old. Bubble size reflects module protein count. NES, normalised enrichment score; WGCNA, weighted gene co-expression network analysis.

Gene ontology enrichment of each module provided biological context (S12 Table; network construction diagnostics in S5 Figure). Per-module z-score heatmaps, eigengene trajectories, and ORA enrichment for the four age- and training-associated modules are shown in S6 Figure. The age-elevated Turquoise module was dominated by organic acid catabolic processes (padj = 2.51 × 10⁻²^0^) and fatty acid catabolism. The training-responsive Blue module was enriched for cell cycle processes including Mitotic Prometaphase (padj = 2.17 × 10⁻⁶). Brown showed strong enrichment for striated muscle contraction (padj = 3.24 × 10^-21^), while Yellow was enriched for glycolysis (padj = 3.11 × 10⁻⁸). Green was dominated by ATP synthesis-coupled electron transport (padj = 2.05 × 10⁻⁵⁷). Black was enriched for translation machinery and Pink for mitochondrial biogenesis.

### Baseline proteomic profiles predict age group and training-induced muscle hypertrophy

To determine which co-expression modules carry outcome-relevant signal, we performed univariate module-by-outcome discrimination using module eigengenes across the annotated WGCNA modules and six outcomes (Fig. 7A). The strongest signals were age-discriminating: Turquoise/M1 (Lipid Catabolism) discriminated Young from Old with AUC = 0.96 (permutation p < 0.001, q < 0.012), followed by Blue/M2 (Cell Cycle/Proteostasis; AUC = 0.89, q < 0.012) and Brown/M3 (muscle contraction; AUC = 0.79, q = 0.077). Within-subject Pre-to-Post timing was weaker across all modules (AUCs 0.46–0.84), and no single module reached BH significance for ΔVL or ΔLBM responder status. Per-module ROC curves across all outcomes are shown in S7 Figure.

**Figure 7.**
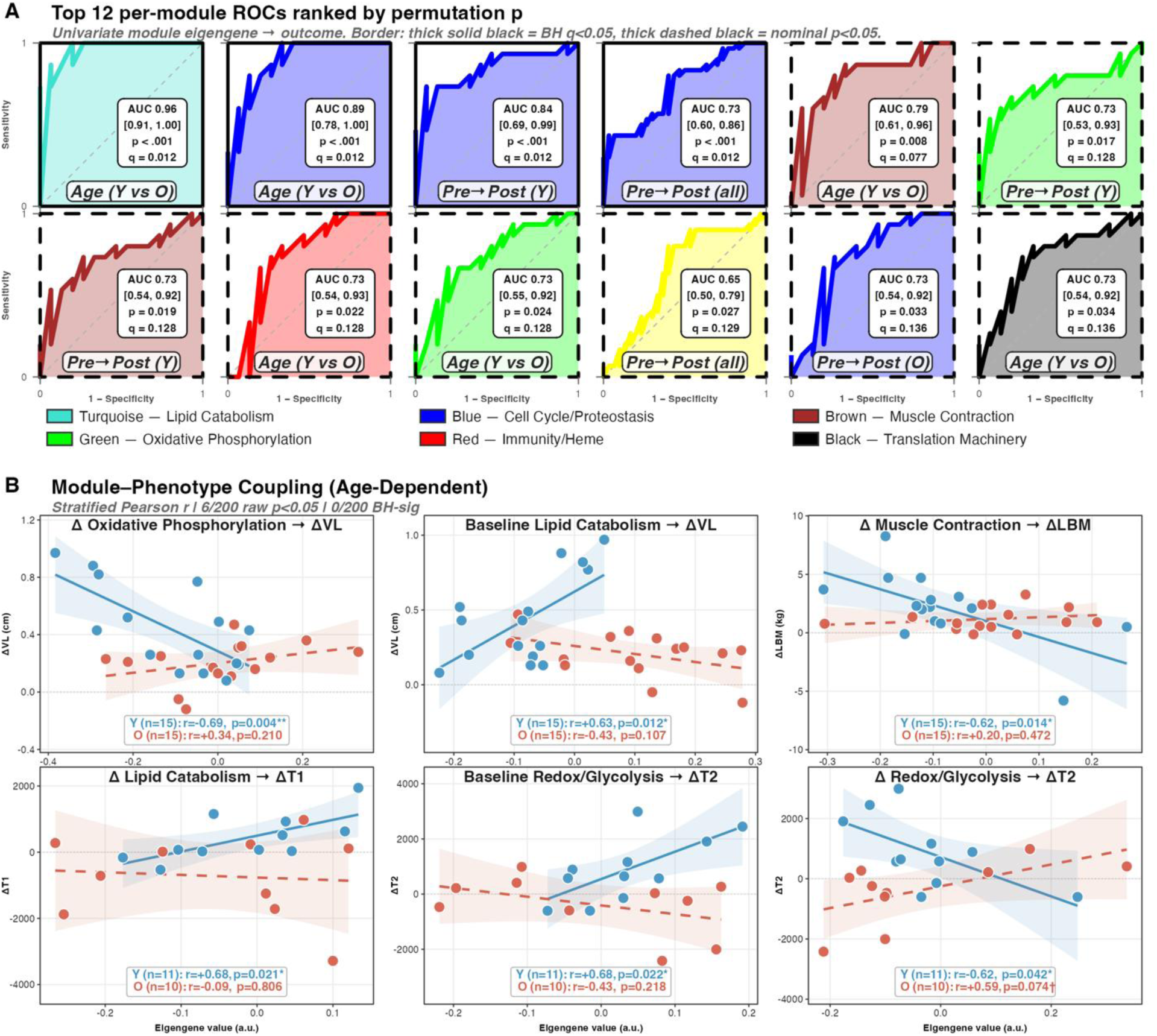
Module eigengene discrimination and age-stratified phenotype coupling. (A) Top-12 univariate ROCs for module eigengenes across six outcomes (Age, Pre→Post timing in Young/All/Old, ΔVL responder, ΔLBM responder). Thick solid border: BH q < 0.05; thick dashed: nominal p < 0.05. Turquoise/Lipid Catabolism (AUC = 0.96) and Blue/Cell Cycle (AUC = 0.89) were the strongest age discriminators (both q < 0.012). (B) Scatter plots of module eigengene values (Δ or baseline) against training-induced phenotypic changes (ΔVL, ΔLBM, ΔType I fCSA, ΔType II fCSA), with stratified Pearson r per age group. Pairs with |r| > 0.50 in at least one stratum are highlighted; none survived BH correction across the 200-test screen. AUC, area under the curve; fCSA, fibre cross-sectional area; LBM, lean body mass; ROC, receiver operating characteristic; VL, vastus lateralis.

Leave-one-subject-out validation confirmed that these in-sample AUCs were not inflated by optimism bias: both fixed-module projection LOSO and full WGCNA-refit LOSO reproduced the Turquoise (0.97 / 0.96), Blue (0.92 / 0.93), and Brown (0.79 / 0.80) out-of-fold AUCs within narrow confidence intervals (S7 Figure). Within-subject change (ΔME) did not discriminate age (AUC = 0.46, p = 0.92), whereas the Pre+Post+ΔME combined stack reached AUC = 0.92 (p = 0.044), supporting the view that age signal lives in static, cross-subject proteome state. Raw-protein (k = 5) Age classifiers achieved AUC = 0.996 (perm p = 0.005), whereas phenotype-only discrimination was not significant (AUC = 0.84, p = 0.12), and smaller WGCNA modules (Red, Yellow, Magenta; Jaccard 0.46–0.66) were less stable under LOSO refit reinforcing that Turquoise, Blue, and Brown carry a coherent age-discriminating signal.

Module eigengenes also tracked phenotypic adaptation with clear age stratification (Fig. 7B). In younger adults, ΔOxidative Phosphorylation was inversely associated with ΔVL (r = - 0.69, p = 0.004), baseline Lipid Catabolism with ΔVL (r = 0.63, p = 0.012), and ΔLipid Catabolism with ΔType I fiber CSA (r = 0.68, p = 0.021). Older adults showed weaker or null associations. These associations were drawn from a 200-test screen (10 modules × 2 sources × 5 outcomes × 2 strata); none survived BH correction across the full screen and should be interpreted as exploratory age-dependent patterns.

Within the age-associated Turquoise module (345 proteins), 30 hub candidates were identified by module membership (kME > 0.75), of which 2 also showed individual association with hypertrophic response (|gene significance for delta VL| > 0.3; ACAT1 |GS| = 0.35, MCCC1

|GS| = 0.32), representing proteins both central to the co-expression network and individually associated with training-induced thickness changes. This provides protein-level evidence that the molecular features distinguishing aging muscle from younger muscle are the same features limiting its capacity for training-induced growth.

## DISCUSSION

The present study leveraged deep quantitative proteomics to characterize how aging modulates the skeletal muscle response to resistance training. Five principal findings emerged. First, despite both age groups implementing a similar 10-12-week program, older adults exhibited markedly attenuated proteome remodeling with 5.6-fold fewer training-responsive proteins and zero FDR-significant proteins. Second, despite this compression, 61.6% of proteins changed in the same direction in both age groups, indicating a conserved but amplitude-limited training response in older individuals. Third, resistance training in older adults partially reversed the aging proteome signature, with ribosomal and translational machinery showing the strongest reversal. Fourth, co-expression network analysis identified age-divergent biological programs, with oxidative phosphorylation emerging as the pathway most resistant to training-mediated reversal and most divergent between age groups. Finally, baseline proteomic profiles were associated with age group membership and training-induced hypertrophic outcomes. Collectively, these results suggest that aging constrains the amplitude of proteome remodeling but does not render the underlying adaptive programs immutable—baseline proteomic state appears to set the boundaries of response capacity, while training can partially restore individual protein abundances within those boundaries. These findings will be expanded upon in the paragraphs below.

### Attenuated but conserved training responses across age

The magnitude of the training response deficit in older adults is substantial. At the Π-score threshold, younger adults produced 100 DEPs compared with 18 in older adults—5.6-fold attenuation (Fig. 2C). At FDR < 0.05, this disparity was more pronounced (134 vs. 0). That the Π-score threshold rescued 18 training-responsive proteins where conventional FDR correction detected none highlights the utility of composite significance metrics that integrate both statistical confidence and biological effect magnitude, particularly in aging populations where genuine but underpowered effects risk being dismissed as null. The consistent pattern across all significance thresholds supports genuine biological attenuation, consistent with O’Leary et al. (32), who reported zero FDR-significant proteins in either age group despite detecting significant pathways, and our past proteomic work showing that aging predominantly affects the non-myofibrillar protein pool with marginal training-induced changes (21). Despite this quantitative deficit, the training response was not qualitatively altered. Protein-level concordance (ρ = 0.284, p = 3.46 × 10⁻⁴⁰; Fig. 4A) showed 61.6% of proteins changing in the same direction in both age groups. RRHO2 analysis confirmed strong concordance with negligible discordant signal (Fig. 4E). This “conservation with compression” pattern suggests that therapeutic strategies aimed at amplifying existing responses, rather than redirecting their trajectory, may hold promise for long-term muscular development in older adults. This is further supported by Lee-Odegard et al. (53), who showed that proteomic aging (ProtAgeGap) was inversely associated with physical activity in 45,438 UK Biobank participants, with a 12-week exercise intervention decreasing ProtAgeGap by approximately 10 months.

Pathway-level concordance (ρ = 0.56, p = 4.40 × 10⁻⁷) substantially exceeded protein-level concordance, suggesting that coordinated functional programs are better preserved across age than individual protein effect sizes, with 23 of 69 curated Hallmark and GO Slim pathways reaching significance in at least one age group (Fig. 4D). The largest age-divergent signal was in oxidative phosphorylation (Hallmark; NES: −2.11 in young vs. −0.62 in old). This extends proteomic observations from transcriptomic precedents such as those by Raue et al. (28) who reported seven-fold fewer responsive transcripts in older adults, and Thalacker-Mercer et al. (30) who reported that baseline transcriptomic profiles predict response heterogeneity. Hence, individual protein effects may be buffered by functional overlap within coordinated pathways, consistent with multi-omic exercise studies (33). In our data, aging and training shared minimal protein-level overlap (85% of Π-score DEPs were contrast-exclusive), yet pathway concordance between age groups (ρ = 0.56) nearly doubled protein concordance (ρ = 0.28). Despite age explaining modest global variance (PERMANOVA R² = 8.9%), 196 DEPs were identified, indicating that aging remodels the proteome through many small, coordinated shifts best captured at the pathway and network level. Notably, Michie et al. (54) reported that older adults exhibit chronically activated unfolded protein response (UPR) pathways at rest, while the adaptive UPR response to acute exercise is similarly activated across age, consistent with the concordant UPR upregulation we observed in both age groups, further underscoring the proteostatic disruptions that characterize aging muscle.

### Resistance training partially reverses the aging proteome

A central finding is that resistance training in older adults directionally opposes age-related proteomic changes. Among 210 proteins meeting the Π < 0.05 threshold across the Aging and Training-in-Old contrasts, 158 (75.2%) showed opposite-sign changes, and among the top 30 aging-affected DEPs, 22 (73%) showed directional reversal (Fig. 5A). RRHO2 revealed asymmetric reversal, with the strongest signal in the Aging-Down/Training-Up quadrant (max −log10p = 35.9; Fig. 5E), extending the transcriptomic reversal reported by Melov et al. (29) to the proteome level. The strongest reversal occurred in ribosomal and translational machinery, with cytoplasmic translation showing complete directional inversion (Aging NES = −2.90; Training NES = +2.60; Fig. 5D). This is particularly notable given evidence that ribosome biogenesis capacity determines hypertrophic outcomes in older adults as mentioned previously. In contrast, oxidative phosphorylation showed minimal reversal, indicating a hierarchy in which translational programs are more responsive to exercise-mediated restoration than mitochondrial metabolic programs. However, a complementary distance-based permutation test applied to the 571 nominally significant aging proteins yielded a non-significant global reversal (11.5%, p = 0.116), indicating that whole-proteome distance-level reversal was modest and not statistically significant at the current sample size.

Our finding that 75.2% of aging-affected proteins show opposite-sign training responses, including 73% of the most strongly affected, suggests that a substantial portion of age-related proteomic change reflects modifiable homeostatic shifts rather than irreversible degeneration.

This is consistent with Hulmi et al. (32), who demonstrated that RT-induced proteomic changes are partially reversible upon detraining, indicating that keletal muscle retains inherent proteome plasticity. Complementary evidence at the epigenomic level further supports this capacity, as DNA methylation signatures of prior hypertrophic stimuli persist through detraining and may prime subsequent retraining responses (47).

### Co-expression network architecture

WGCNA identified ten co-expression modules organizing 1,606 of 2,113 proteins (507 unassigned). Using linear mixed models to account for the paired design, three modules were significantly associated with aging: Turquoise (lipid catabolism; r = +0.59, pBH = 4.6 × 10⁻⁵), Magenta (translation initiation; r = −0.53, pBH = 5.7 × 10⁻⁴), and Brown (muscle contraction; r = −0.44, pBH = 0.008). Blue (cell cycle/proteostasis; r = +0.63, pBH = 0.002) was the only module significantly associated with training in younger adults, suggesting these programs are preferentially reactivated by RT. The Green module (oxidative phosphorylation) eigengene change was inversely correlated with hypertrophy in younger adults (r = −0.69, pBH = 0.046), reinforcing the pathway-level OxPhos suppression observed in this group. Black was enriched for translation machinery—aligning with the ribosomal divergence observed in the concordance and reversal analyses. Nine of ten modules were strongly preserved between pre- and post-training networks (Z > 10), with Purple showing moderate preservation (Z = 8.6). Hence, the co-expression architecture of the skeletal muscle proteome is largely stable across the training intervention, even as individual protein abundances shift. This modular organization parallels WGCNA-derived transcriptional networks identified by Lavin et al. (55), who showed that baseline molecular network state shapes hypertrophic response heterogeneity in older adults undergoing 14 weeks of RT, further supporting that network-level proteome architecture constrains adaptive capacity. Notably, these module-level findings recapitulate the reversal hierarchy observed at the pathway level: translation-associated modules (Magenta, Black) align with the programs most responsive to training-mediated reversal, whereas the Green/oxidative phosphorylation module, whose lack of response tracked with reduced hypertrophy, corresponds to the pathways least amenable to exercise-induced restoration. This suggests that the same biological programs resistant to reversal are those whose persistent dysregulation limits adaptive outcomes.

### Mitochondrial and metabolic pathway remodeling

The pronounced age-dependent divergence in mitochondrial pathway responses warrants discussion. Younger adults showed strong suppression of oxidative phosphorylation following training (Hallmark OxPhos NES = −2.11), which may reflect a shift in biosynthetic resources toward growth-related programs, while this response was almost entirely absent in older adults (NES = −0.62), generating the strongest interaction effect among all pathways (NES = +2.02). Aging itself was associated with elevated oxidative phosphorylation gene products (NES = +2.00), which may represent compensatory upregulation in response to declining mitochondrial efficiency. Robinson et al. (35) reported that resistance training increased mitochondrial protein synthesis in older but not younger adults, suggesting older muscle responds to energy demands by augmenting mitochondrial biogenesis rather than undergoing the glycolytic shift observed in younger adults, though Jessen et al. (56) recently showed that RT-induced OxPhos suppression occurs predominantly in type I fibers, suggesting the whole-muscle signal reflects fiber-type-specific remodeling. This age-specific failure to remodel mitochondrial protein content may reflect impaired mitochondrial quality control, as mitochondrial dysfunction is increasingly recognized as both a hallmark and potential driver of biological aging. The MoTrPAC mitochondrial atlas further demonstrates tissue-specific mitochondrial multi-omic remodeling during endurance training (57), with skeletal muscle showing an absence of a coordinated oxidative protein upregulation in our RT context.

### Baseline proteomic signatures predict age and hypertrophic adaptation

Module eigengenes decomposed age- and training-related signal across distinct biological programs. Univariate per-module ROCs indicated that age separation is carried primarily by metabolism-linked modules (Turquoise/Lipid Catabolism, AUC = 0.96, q < 0.012) and proteostasis (Blue/Cell Cycle, AUC = 0.89, q < 0.012; Fig. 7A), with Brown/Muscle Contraction providing a weaker nominal signal (AUC = 0.79, q = 0.077), while within-subject training-time discrimination was weaker across all modules (AUCs 0.65–0.79). Module eigengenes also tracked phenotypic adaptation with clear age stratification (Fig. 7B): in younger adults, ΔOxidative Phosphorylation was inversely associated with ΔVL (r = -0.69, p = 0.004), baseline Lipid Catabolism tracked ΔVL (r = 0.63, p = 0.012), and ΔLipid Catabolism tracked ΔType I fiber CSA (r = 0.68, p = 0.021), while older adults showed weaker or null associations. Within the age-associated Turquoise module (345 proteins), 30 hub candidates (kME > 0.75) showed a consistent anti-correlation between age-associated and hypertrophy-associated proteins, providing protein-level evidence that the molecular features distinguishing aging muscle directly limit its growth potential.

### Mechanistic and clinical implications

The convergence of multiple independent analytical approaches—differential expression, concordance, and network analysis—on common biological themes insights into how aging can affect the proteome. The pronounced attenuation at the protein level, combined with preserved pathway-level concordance, suggests that aging introduces bottlenecks at transcriptional and post-transcriptional levels that compress individual protein effect sizes while preserving functional program coordination. Hence, we posit that the anabolic signaling infrastructure necessary for maximal proteome remodeling is selectively impaired with aging. The conservation-with-compression pattern implies that therapeutic strategies should aim to amplify existing response amplitudes rather than redirect response trajectories. The ribosome biogenesis pathway emerges as a particularly attractive intervention target given that ribosomal proteins were among the most strongly restored by training in older adults (Figs. 5C–D), and ribosome biogenesis capacity predicts hypertrophic outcomes as previously stated; though the relative importance of translational capacity versus efficiency may shift as training progresses, with greater hypertrophy linked to lower ribosome-related gene expression after prolonged RT (58). While speculative, pharmacological, nutritional, or other interventional strategies aimed at enhancing both ribosome biogenesis and efficiency—such as leucine supplementation or resistance exercise combined with essential amino acid provision—may augment the amplitude of proteome remodeling in older adults.

### Limitations

This study has several limitations. First, the analysis is secondary, drawing from parent RT investigations with shared training protocols but different nutritional supplementation paradigms (peanut protein and beetroot juice), and participant-level random intercepts accounted for within-subject correlation. Second, the sample sizes (n = 17 younger, n = 15 older) provided adequate power for detecting moderate-to-large training effects in younger adults but limited power for the interaction contrast. The complete absence of FDR-significant findings in Training Old likely reflects both biological attenuation and power limitations. Third, the aging component is cross-sectional, and comparisons between age groups may reflect cohort differences rather than biological aging per se, and a single post-training timepoint cannot capture the kinetics of proteome remodeling. Fourth, all participants were male, and results may not generalize to females given known sex differences in skeletal muscle physiology and training adaptations. Additionally, all biopsies were obtained from the VL, and results may not generalize to muscles with different fiber-type compositions or functional demands, particularly upper-body or predominantly slow-twitch muscles such as the soleus. Age-related impairments in recovery from exercise-induced muscle damage (59) may also produce differential adaptive responses depending on the RT protocol employed, and results be interpreted within the context of the specific training design used herein. Finally, the older cohort (mean age ∼58 years) is younger than many participants sampled in aging studies across the literature, and results may underestimate effects in adults over 65 years old.

## Conclusions

In summary, these data provide evidence that aging markedly attenuates but does not qualitatively alter skeletal muscle proteome plasticity in response to resistance training. The training response in older adults, while compressed in magnitude, retains directional conservation with younger adults and directionally opposes a substantial fraction of age-related proteomic changes. Ribosomal and translational machinery are the most responsive to training-mediated reversal, while mitochondrial metabolic programs are the least. Critically, given the lack of age-related proteome comparisons to resistance training, future research is needed to provide continued insight into this area of skeletal muscle biology.

## ACKNOWLEDGMENTS

We thank the participants who devoted time to volunteer for the studies used for this analysis. We also thank past laboratory members who assisted with the prior studies.

## DATA AVAILABILITY

Processed proteomics data will be deposited in a public repository (PRIDE/MassIVE) upon publication. Analysis code is available at https://github.com/Dustyn-T-Lewis/YvO_2025.

## AUTHOR CONTRIBUTIONS

DTL, ANK, and MDR conceived the idea for the manuscript. JMM, MCM, DRT, DLP, MLM, NJK, GK Jr, BJM, and JSG assisted with several aspects of the parent studies which warrant co-authorship herein. SCN and MDB were involved in proteomics analysis and intellectual input during data processing. ADF and CBM provided significant supervision of students during the execution of the three studies. All authors provided intellectual insight and/or edits to the manuscript, and all authors approved the final submitted version.

## SUPPLEMENTAL APPENDIX

S1 Table. Per-sample metadata and baseline phenotype for the analyzed cohort (64 samples; 62 retained, 2 excluded; QC_Status column identifies consensus outliers).

S2 Table. Raw DIA-MS protein intensity matrix prior to normalization (2,837 proteins × 64 samples–before exclusion).

S3 Table. Cyclic-loess–normalized log₂ protein matrix with per-protein and per-sample missingness summaries (2,113 proteins × 62 samples).

S4 Table. Post-imputation protein matrix (missForest; ranked second in a 17-method benchmark, selected for superior fold-change preservation and parsimony; OOB NRMSE = 0.109) with Boolean imputation mask and per-protein imputation check.

S5 Table. Imputation benchmark (17-method composite ranking and per-method metrics) and per-protein MAR vs MNAR classification (3-method consensus; missForest treats all NAs as MAR).

S6 Table. Differential expression results across four contrasts with Π-score (per Xiao et al. 2014), FDR, and post hoc power analysis.

S7 Table. Figure 1 source data — phenotypic responses to resistance training (training volume, lean body mass, VL thickness, baseline characteristics) and S2 Figure panel data.

S8 Table. Figure 2 source data — proteome overview and DEP landscape (PCA, log₂FC density, DEP counts, UpSet intersections, fGSEA, rank barcode) and S3 Figure panel data.

S9 Table. Figure 3 source data — volcano ring plots, per-contrast pathway enrichment, and S4 Figure panel data (model diagnostics).

S10 Table. Figure 4 source data — training concordance between younger and older adults (ORA quadrants, NES scatter, pattern heatmap, fry, RRHO2) and S8 Figure panel data (ORA dedup sensitivity, ρ bootstrap, threshold sensitivity, GO Slim distribution, fry leading edge).

S11 Table. Figure 5 source data — aging reversal analyses (ORA quadrants, NES scatter, pattern heatmap, fry, RRHO2; includes per-protein reversal/exacerbated/negligible directional classification).

S11 Table. Figure 5 source data — aging reversal analyses (ORA quadrants, NES scatter, pattern heatmap, fry, RRHO2; includes per-protein reversal/exacerbated/negligible directional classification) and S9 Figure panel data (ORA dedup sensitivity, r bootstrap, circularity diagnostic, threshold sensitivity, GO Slim distribution, fry leading edge).

S12 Table. Figure 6 source data — WGCNA module assignments, hub proteins (kME ≥ Q90), module enrichment, module-trait associations (LMM with Kenward-Roger df, BH-corrected), triptych enrichment and module eigengenes.

S13 Table. Figure 7 source data — univariate per-module ROCs, leave-one-subject-out validation, and age-stratified module-phenotype coupling.

S1 Figure. Quality-control pipeline summary. (A) Protein filter cascade (2,837 to 2,113). (B) Per-sample missingness. (C) Pre-normalization PCA. (D) Post-normalization PCA. (E) Missingness classification. (F) Imputation effect density (OOB NRMSE = 0.109).

S2 Figure. Supplementary phenotype responses. (A) Deadlift one-repetition maximum. (B) Type II fiber cross-sectional area. (C) Type I fiber cross-sectional area.

S3 Figure. Proteome-level coefficient of variation by experimental group.

S4 Figure. Differential expression model diagnostics across contrasts. (A) P-value histograms. (B) Π-score distributions. (C) FDR distributions. (D) MA plots. (E) Imputation sensitivity. (F) Outlier sensitivity.

S5 Figure. WGCNA network construction diagnostics. (A) Scale-free topology fit. (B) Protein dendrogram with module assignments. (C) Subcellular compartment enrichment. (D) Pearson vs. biweight midcorrelation module overlap.

S6 Figure. Module triptychs for four key co-expression modules (Turquoise/Lipid Catabolism, Magenta/Translation Initiation, Brown/Muscle Contraction, Blue/Cell Cycle). Each row: z-score heatmap, eigengene trajectories, and ORA enrichment.

S7 Figure. Module eigengene discrimination and cross-validation. (A) Per-module ROC grid across all module-outcome combinations. (B) Fixed-module leave-one-subject-out sensitivity. (C) Full WGCNA-refit leave-one-subject-out sensitivity.

S8 Figure. Training concordance robustness diagnostics. (A) ORA pathway counts across Jaccard deduplication cutoffs. (B) Spearman ρ bootstrap (1,000 replicates). (C) Quadrant counts across significance thresholds. (D) GO Slim category distribution by quadrant. (E) Top 20 fry leading-edge proteins in Training Old.

S9 Figure. Aging reversal diagnostic panels. (A) ORA deduplication sensitivity for reversal quadrants (Jaccard cutoffs 0.3–1.0). (B) Pearson r bootstrap (1,000 replicates, 95% CI) for Aging vs. Training-in-Old log₂FC correlation. (C) Circularity diagnostic: protein-permuted null distribution of Pearson r (observed r = −0.358, null mean = 0.0008, p_perm < 0.001). (D) Reversal classification threshold sensitivity (stable reversal percentage across |log₂FC| thresholds 0.05–0.30). (E) GO Slim category distribution of reversed and non-reversed proteins (210 DEPs across 15 categories). (F) fry leading-edge proteins: top 25 reversal drivers ranked by |t-statistic| in Training Old, colored by aging direction.

